# The BNT162b2 mRNA vaccine induces polyfunctional T cell responses with features of longevity

**DOI:** 10.1101/2021.09.27.462006

**Authors:** Gisella Guerrera, Mario Picozza, Silvia D’Orso, Roberta Placido, Marta Pirronello, Alice Verdiani, Andrea Termine, Carlo Fabrizio, Flavia Giannessi, Manolo Sambucci, Maria Pia Balice, Carlo Caltagirone, Antonino Salvia, Angelo Rossini, Luca Battistini, Giovanna Borsellino

**Affiliations:** Neuroimmunology Unit, Santa Lucia Foundation IRCCS; Rome, Italy; Data Science Unit, Santa Lucia Foundation IRCCS; Rome, Italy; Clinical Microbiology Laboratory, Santa Lucia Foundation IRCCS; Rome, Italy; Department of Clinical and Behavioral Neurology, Santa Lucia Foundation IRCCS; Rome, Italy; Medical Services, Santa Lucia Foundation IRCCS; Rome, Italy

## Abstract

Vaccination against SARS-CoV-2 infection has shown to be effective in preventing hospitalization for severe COVID-19. However, multiple reports of break-through infections and of waning antibody titers have raised concerns on the durability of the vaccine, and current discussions on vaccination strategies are centered on evaluating the opportunity of a third dose administration. Here, we monitored T cell responses to the Spike protein of SARS-CoV-2 in 71 healthy donors vaccinated with the Pfizer–BioNTech mRNA vaccine (BNT162b2) for up to 6 months after vaccination. We find that vaccination induces the development of a sustained anti-viral memory T cell response which includes both the CD4+ and the CD8+ lymphocyte subsets. These lymphocytes display markers of polyfunctionality, are fit for interaction with cognate cells, show features of memory stemness, and survive in significant numbers the physiological contraction of the immune response. Collectively, this data shows that vaccination with BNT162b2 elicits an immunologically competent and potentially long-lived SARS-CoV-2-specific T cell population. Understanding the immune responses to BNT162b2 provides insights on the immunological basis of the clinical efficacy of the current vaccination campaign and may instruct future vaccination strategies.

## Introduction

Mass-vaccination against COVID-19 has quickly shown to be effective and to confer high levels of protection against COVID-19 in real-world settings *(1)*. However, one notion that all immunologists learned during the pandemic was that natural infection with coronaviruses induces short-lived immunity *(2)*, with reinfections occurring frequently. This notion was quickly revised by subsequent studies showing that T cell immunity to SARS is actually long lived *(3, 4)*. The study of immune responses in COVID-19 patients with various degrees of disease severity have revealed that indeed infection with SARS-CoV-2 elicits a robust immune response, involving both the innate and the adaptive arms of the immune system and theoretically effective in protecting from reinfections *(5, 6)*. However, this protection is not absolute and impeccable, and cases of reinfection do occur *(7)*. Humoral immunity provides a shield against reinfection through the generation of neutralizing antibodies, which are easily measurable and are widely used as indicators of protective immunity *(8, 9)*. Notably, subjects with undetectable or impaired humoral responses can nonetheless clear the infection, suggesting that antigen-specific T cells are themselves effective at containing the virus *(10–12)*. Evidence of T cell involvement in the immune response to SARS-CoV-2 came from reports showing that the emergence of activated T cells precedes recovery from COVID-19 *(13–16)*, and was later confirmed in *in vitro* studies which characterized antigen-specific CD4+ and CD8+ lymphocytes reactive with overlapping peptide pools from the SARS-CoV-2 Spike protein *(15, 17–19)*. These cells persist in COVID-19 convalescents *(4, 20–23)*, and have been shown to reduce viral loads in non-human primate models of infection; crucially, they arise also following vaccination both with mRNA- and adenoviral-vaccines *(24–27)*. It is necessary and urgent to identify the cellular immune components induced by vaccination and to measure their persistence, also in the view of reports of waning immunity over time and in recent updates in vaccination strategies which now propose administration of a third dose. Here, we show the results of a longitudinal study on T cell responses in 71 health-care workers and scientists vaccinated with the BNT162b2 vaccine following the European Medicines Agency (EMA)-approved vaccination schedule, up to 6 months after the first dose. We find robust induction of neutralizing antibodies and of Spike-specific CD4+ and CD8+ lymphocytes which persist in the periphery after the physiological contraction of the initial response. These cells are detectable in most individuals also before vaccination, denoting the presence of a pre-existing pool of cross-reactive cells, and they are expanded significantly after the boost. We show that vaccine-induced T cells are mostly central and effector memory cells, and that they are equipped with overlapping sets of molecules which enable them to perform multiple immune functions: facilitation of B cell differentiation and antibody production, direct cytotoxic activity, and cytokine production, and that they are equally represented in both sexes. Importantly, we show that vaccination also induces the generation of potentially long-lived memory stem cells, pilasters of durable immunity.

## Results

### Induction and persistence of neutralizing antibodies

The antibody response to vaccination with BNT162b2 was measured in serum samples obtained at the day of the boost (T1), 14 days later (T2), and 6 months after the first dose (T3). As expected from previous studies, all individuals in our cohort were devoid of neutralizing antibodies (nAbs) at baseline, and significant levels of nAbs were obtained in 100% of individuals only after the second dose (Fig.1A), (T1: median 28; T2, median 1786; T3: median 517). Age and gender are key variables in immunity induced by vaccination, whose effectiveness decreases with age and is usually lower in males *(28)*. Thus, we analyzed the differences in antibody levels in male and female donors and also correlated them with age. We find that antibody titers correlate negatively with age at all time points, confirming previous results *(29, 30)* (Fig. 1C). In our cohort antibody levels induced by BNT162b2 are comparable between males and females, as shown also in other studies *(30)*(Fig. 1B), although females seem to retain lower levels of nAbs 6 months after vaccination. Thus, this data shows that BNT162b2 aptly induces the production of nAbs which decrease over time but are however maintained at high levels for at least 6 months.

**Fig. 1:**
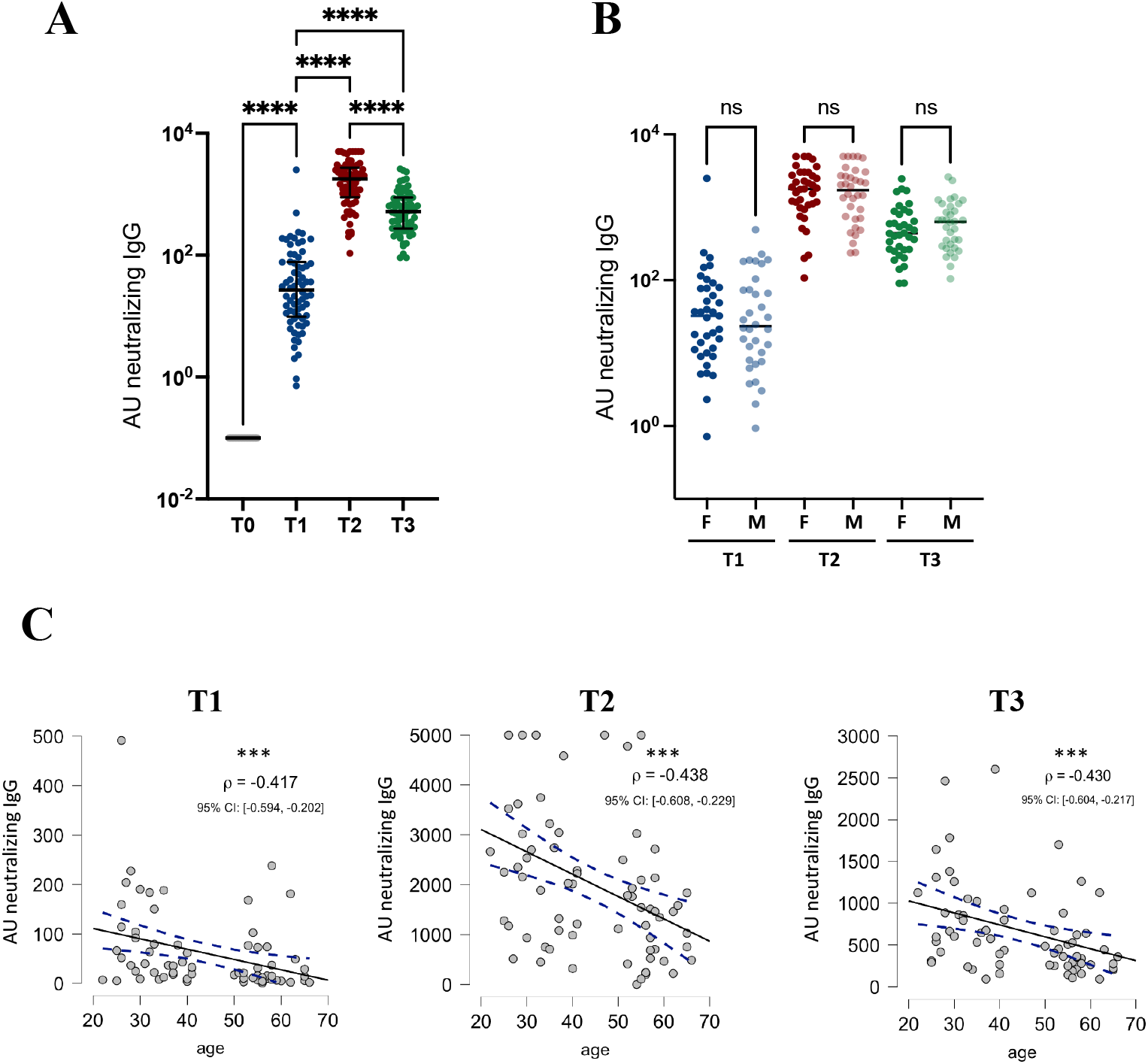
Antibody response following vaccination with BNT162b2. A) Concentration of neutralizing anti-Spike IgG at baseline (T0), 21 days after the first dose (T1), 14 days after the second dose (T2), and 6 months after the first dose (T3). p values were determined using the Friedman test with Dunn correction for multiple comparisons; ***p < 0.001. **** p < 0.0001; no simbols: not significant. B) antibody levels at the different time points in females (F) and males (M). Statistical analysis between time points was performed using Kruskall-Wallis test. ns= not significant C) Spearman’s rank correlation test was applied to test for correlation between AU neutralizing IgG and age at each time point (T). Significance levels, Spearman’s rank correlation coefficient (ρ) and Confidence Intervals for each plot are reported in figure (* = p < 0.05; ** = p < 0.01; *** = p < 0.001; **** = p < 0.0001).

### Induction and durability of the Spike-specific T cell response

To investigate the cellular immune response induced by vaccination, we exposed freshly obtained peripheral blood mononuclear cells (PBMC) to peptide pools spanning the entire sequence of the Spike (S) protein. Blood collection was performed at baseline (T0), on the day of the boost (day 21, T1), 14 days later (T2), and after 6 months (T3). Several effector functions of CD4+ cells consist in the upregulation of surface molecules for intercellular communication, and these may remain trapped inside the cells if the secretion inhibitors required for intracellular cytokine detection are present during antigenic stimulation. Thus, to fully capture the T cell antigen-specific response and to maximize sensitivity, two separate assays for the detection of the expression of surface Activation-Induced Markers (AIM) and for Intracellular Cytokine Staining (ICS) were set up (Fig. S1 for gating strategy). Moreover, all assays were performed on freshly isolated PBMCs in order to avoid the inevitable cell loss during the freezing/thawing procedure, and to obtain accurate absolute cell counts.

AIM+ CD4+ T cells were defined by upregulation of CD40L and CD69, while CD137 and CD69 expression identified the AIM+ CD8+ subset (Fig. 2A). After paired background subtraction from parallel unstimulated cultures, 99% of donors (66/67) had detectable numbers of AIM+ CD4+ cells at baseline (median 349 cells/ml, range 63-4639); 21 days after the first dose of vaccine these cells were identified in all donors, and at higher levels (median 2218 cells/ml, range 319-110394, p <0.0001), and further still at 14 days after the boost (median 3608 cells/ml, range 566-101864); 6 months after the first dose AIM+ CD4+ cell numbers were still increased 5-fold compared to baseline (median 1660, range 349-74859). AIM+ CD8+ T lymphocytes were present at baseline in only 63% of the donors (42/67, median 209 cells/ml, range 18-2695), and increased following vaccination (59/71, median 900 cells/ml, range 54-21142, p<0,0001), with further expansion after the second dose (64/69, median 1539 cells/ml, range 46-18,638), and persisted after 6 months in all donors (median 706 cell/ml, range 78-27593) (Fig. 2B,C).The total magnitude of the T cell response (that is, including both CD4+ and CD8+ cells) increased significantly following priming (T1) and rechallenge (T2), and decreased 6 months after the first dose (T3) (Fig. 2D). Importantly, the Stimulation Index (SI, the ratio of AIM+ T cells in stimulated over unstimulated samples) of AIM+ CD4+ cells increased dramatically after the first dose, remained at high levels after boosting (Fig. 2E), and increased further after 6 months, reaching a median of 25,7. The SI of AIM+ CD8+ T cells peaked 14 days after the boost, and declined by 1/3 at the latest time-point. Moreover, the fraction of individuals showing CD4+ with SI >3 increased at all time points, denoting the establishment of an antigen-specific T cell population (Fig. 2F).

**Fig.2:**
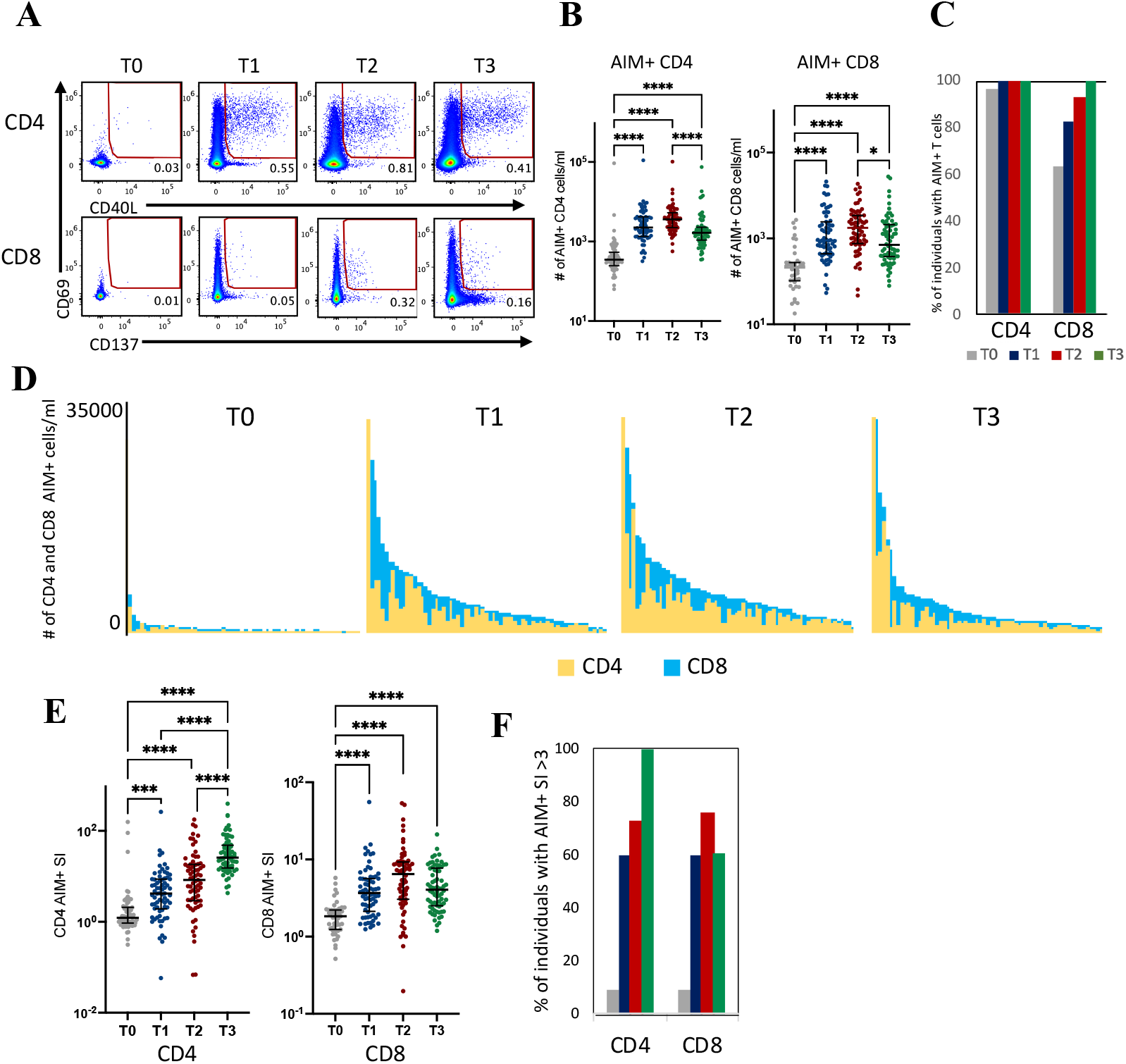
Spike-specific T cell responses induced by vaccination with BNTb16b2. A) Representative flow cytometry plots gated on CD4+ or CD8+ T cells showing upregulation of activation markers (CD69 and CD40L for CD4 cells and CD69 and CD137 for CD8 cells) following o.n. stimulation with a pool of overlapping peptides covering the wt Spike protein at baseline (T0), 21 days after the first dose (T1), 14 days after the second dose (T2), and 6 months after initial vaccination (T3). Numbers in gates represent percentages of positive cells. B) Longitudinal analysis of spike-specific CD4+ and CD8+ absolute cell counts, following background subtraction, in paired samples. Time points were compared by non parametric Kruskall Wallis repeated measures Friedman test; lines represent median with interquartile range. *p < 0.05; **p < 0.01; *** p < 0.001; ****p < 0.0001; no symbol, not significant. C) Fraction of individuals showing spike-specific AIM+ CD4+ and CD8+ cells at each time point. D) Magnitude of T cell responses. AIM+ CD4+ and CD8+ absolute cell counts were aggregated for each donor and displayed in the bar charts at the different time points. E) Stimulation indeces (ratio of stimulated and paired unstimulated samples) of CD4 and CD8 cells in the different time points; time points were compared by non parametric Kruskall-Wallis test; bars represent median with interquartile range. *p < 0.05; **p < 0.01; *** p < 0.001; ****p < 0.0001; no symbol, not significant. F) Fraction of individuals showing S.I. >3 in CD4 and CD8 cells at the different time points. G)

The frequency of S-specific CD4+ (and not CD8+) T cells induced by vaccination correlated inversely with age only 21 days after the first dose (T1), although a tendency towards reduced numbers of activated T cells with increasing age was observed at all time points (Fig. S2A), again with no significant differences between males and females (Fig. S2B).

The numbers of AIM+ cells correlated with antibody levels only after priming and not at subsequent time points (Fig. S2C), suggesting that humoral and cellular immune responses follow different kinetics. However, machine learning indicated that the number of AIM+ CD4+ cells 21 days after priming is the best predictor of nAbs levels after 6 months, as expected from a T cell-dependent B cell response (Fig. S2D).

Thus, vaccination induces clearly detectable and robust antigen-specific T cells in both sexes, arising before the development of high antibody titres, with a major expansion occurring after the first dose of vaccination followed by consolidation of the response after the boost, and persistence for up to 6 months.

### Cytokine production by S-specific T cells

Cytokine production was assessed by intracellular flow cytometry following stimulation o.n. with the peptide pools in the presence of monensin and brefeldin (Fig. 3A). Measurement of cytokine production showed that at baseline 62% and 50% of the tested donors showed IFNγ+ CD4+ and CD8+ cells, respectively (CD4: median 90 cells/ml, range 1-1536; CD8: median 116 cells/ml, range 9-592) (Fig. 3B,C). Three weeks after the first dose, 90% of individuals showed IFNγ+ CD4+ T cells (median 275 cells/ml, range 5-5799), while CD8+IFNγ+ (median 446 cell/ml, range 2-6606) were found in 77% of individuals. Two weeks after the boost, 97% and 69% of individuals showed IFNγ+ CD4+ (median 17111 cell/ml, range 200-19595) and CD8+ T cells (median 592 cells/ml,, range 1-36917), respectively; these cells were maintained for 6 months (CD4: median 360 cell/ml, range 51-13028; CD8: median 315, range 13-4586). 53% of donors at baseline showed IL-2+ CD4+ T cells (median 13 cells/ml, range 2-171); this fraction increased to 84% after the first dose (median 139 cells/ml, range 4-2595) then reached 94% after the boost (median 1838 cells/ml, range 140-17302), and remained detectable after 6 months (median 446 cells/ml, range 20-5678). Polyfunctional IFN-γ+IL-2+ CD4+ T cells were induced in 99% of the individuals only following the booster dose (median 825 cells/ml, range 43-7088), and persisted for at least 6 months after the first dose (median 169 cells/ml, range 9-1533) (Fig. 3D). Polyfunctional CD8+ T cells co-expressing IFNγ+ and lysosomal associated membrane glycoprotein (LAMP-1, CD107a) increased significantly after the boost (T0: median 33 cells/ml, range 1-1528; T1: median 54 cells/ml, range 2-694; T2: median 108 cells/ml, range 11-11965; T3: median 106 cells/ml, range 1-1854) (Fig. 3D). Induction of these cytokine-secreting S-specific T cells was equivalent in both sexes at all time points (Fig. S3A). The number of TNF-α-secreting CD4+ and CD8+ cells was highest at the peak of the secondary response, 14 days after the second dose (Fig. S3B); at the same time point polyfunctional TNFα+IFNγ+ cells decreased significantly, and after 6 months they were found to be at pre-vaccination levels. Further analysis of CD107a, TNFα and Granzyme B co-expression among CD4+ and CD8+ T cells producing IFNγ+ revealed changing patterns between time points (p<0.001 for all comparisons except for IFNγ+ CD4+ T cells between T1 and T2, Figure 3E), with the fraction of polyfunctional CD4+ cells increasing at each time point, and a predominance of CD8+ cells with 2 functions at the latest time point, indicating a dynamic refolding of functional programs in antigen-specific T cells following boost and and contraction.

**Fig.3:**
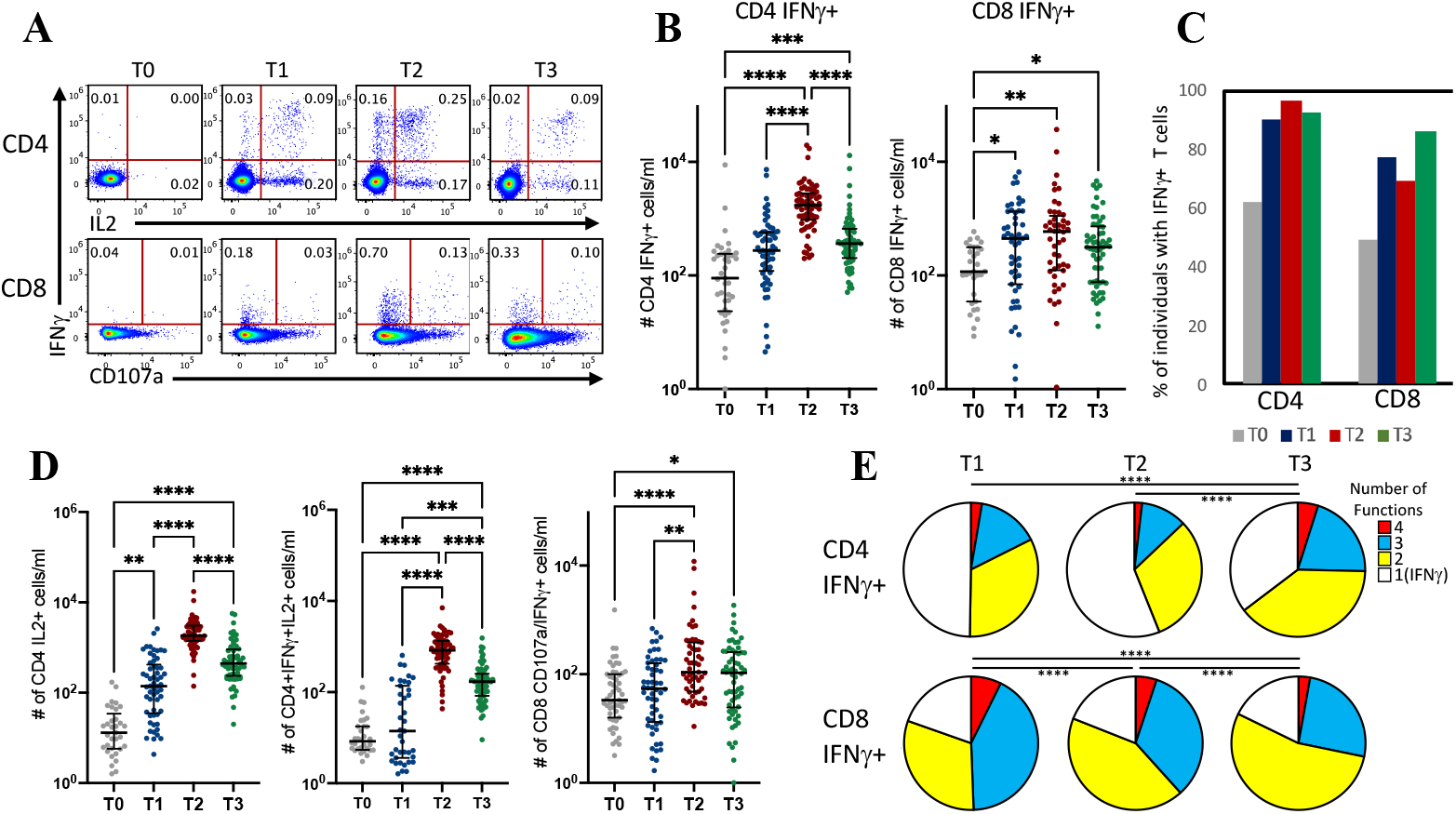
Spike-specific T cell responses are characterized by cytokine production and by markers of cytotoxicity. A) Representative flow cytometry plots gated on CD4+ or CD8+ T cells showing production of IFNγ and IL2 by CD4+ cells (top) and both IFNγ production and upregulation of CD107a (LAMP-1) by CD8 cells (bottom) following stimulation with a pool of overlapping peptides covering the wt Spike protein, at baseline (T0), 21 days after the first dose (T1), 14 days after the second dose (T2), and 6 months after initial vaccination (T3). Numbers in gates indicate percentages of positive cells. B) Longitudinal analysis of absolute cell counts of spike-specific IFNγ producing CD4+ and CD8+, following unstimulated control background subtraction, in paired samples. Time points were compared by non parametric repeated measures Friedman test; lines represent median with interquartile range. *p < 0.05; **p < 0.01; *** p < 0.001; ****p < 0.0001; no symbol, not significant. C) Fraction of individuals showing spike-specific IFNγ+ CD4+ and CD8+ cells at each time point. D) Longitudinal analysis of Spike-specific CD4+ cells producing IL2 or both IL2 and IFNγ (left and center panels) and of CD8+ cells expressing both CD107a and IFNγ (left panel). p values were determined as in Fig 3B. E) Coexpression of functional markers (CD107a, Granzyme B and TNFα) in IFNγ-producing S-specific CD4+ and CD8+ T cells. The percentage of T cells positive for the specified number of functions is indicated by the pie slices for each timepoint, with functions=0 indicating the fraction of cells that produce only IFNγ (gating parameter). ****: p < 0.001 by a partial permutation test (10000 iterations, Monte Carlo simulation) on distributions into combinatorial gates.

Cytokine secretion was also measured, at the peak of the response (T2), in the supernatants of cultures stimulated with the peptide pools. Production of high levels of IFNγ and IL-2 was confirmed, whereas IL-17 and IL-4 were barely detectable (<5 pg/ml), confirming the Th1 differentiation profile of S-specific cells (Fig. S3B).

Thus, vaccination induces the emergence in both males and females of a robust CD4+ and CD8+ cytokine response by T cells already after priming, while full effector functions marked by polyfunctionality are acquired only following the boost, and then maintained for at least 6 months.

### Features of Spike-specific T cells

We then characterized the antigen-specific T cells induced by vaccination through the definition of their differentiation status (Fig.4). The study of the composition of naïve, memory, and effector cell fractions within AIM+ cells in the peripheral blood shows a high frequency of cells with a naïve phenotype at baseline which drops after vaccination, matched by a striking increase in the fraction of effector and central memory cells in both CD4+ and CD8+ subsets, suggesting differentiation from a pool of S-specific cell precursors after antigen exposure and successful induction of a memory cell pool (Fig. 4A and B). The fraction of terminally differentiated CD45RA+CCR7- in both CD4+ and CD8+ subsets drastically decreases following vaccination, suggesting the establishment of a lively and non-terminal immune response. Six months after the first dose, S-specific cells which have survived the physiological contraction of the immune response are mostly central- and effector memory cells, and in the CD8+ subset these cells also include a significant fraction of terminally differentiated effectors. It should be noted that the fraction of AIM+ T cells at baseline necessarily includes also non Spike-specific CD4+ and CD8+ lymphocytes, (as suggested by the low stimulation index and by the similar distribution of the differentiation subsets in AIM+ and in total CD4+ or CD8+ T cells, respectively), and these cannot be excluded from the analysis. After priming and boosting, however, the very high stimulation indices indicate that AIM+ cells are completely represented by S-specific T lymphocytes. For this reason, we limited further and deeper phenotypic analysis of Spike-specific cells to the time points after vaccination.

**Fig.4:**
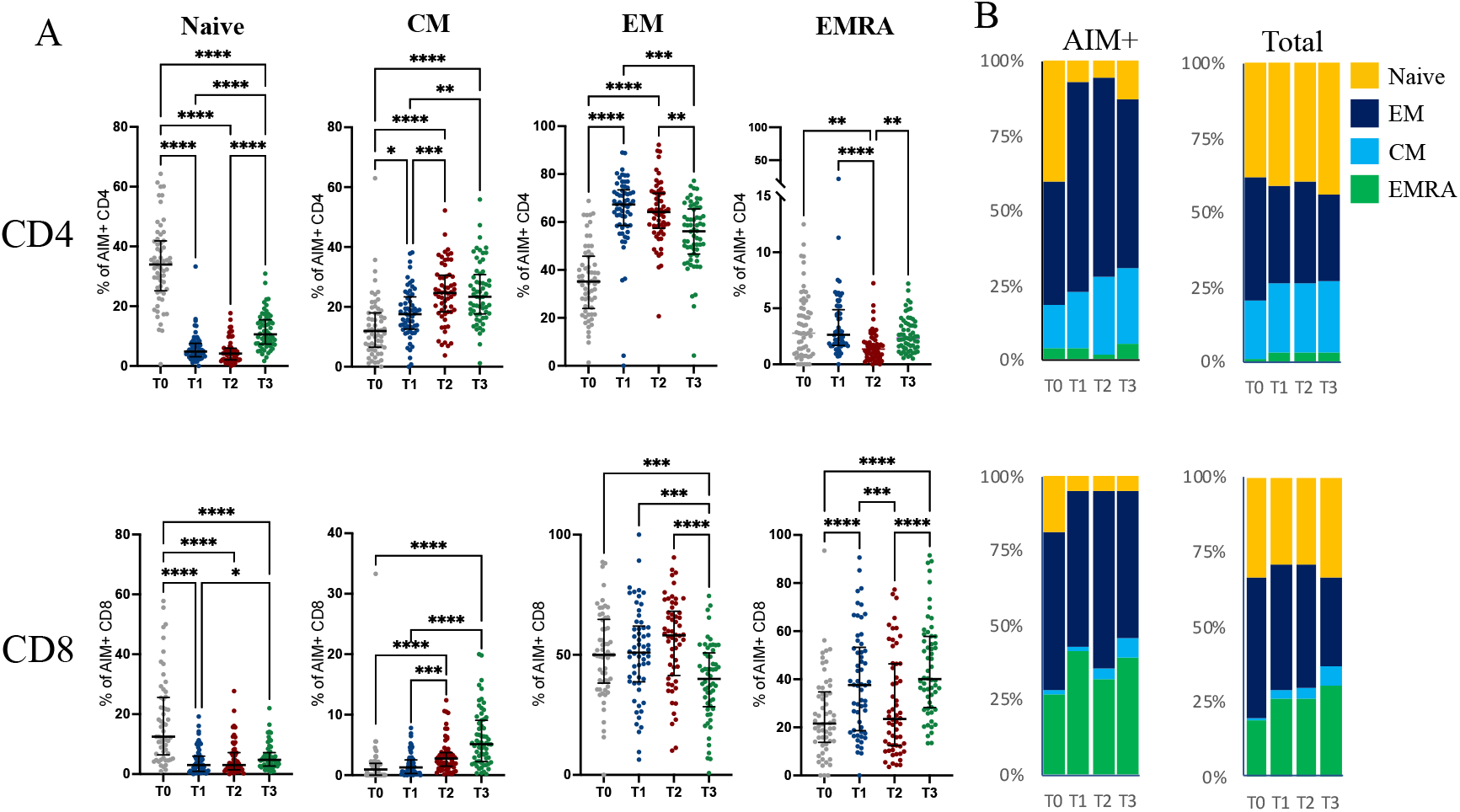
Differentiation status of Spike-specific CD4+ and CD8+ T cells. A) Frequency of naïve, central memory (CM), effector memory (EM) and CD45RA+ effector memory (EMRA) within AIM+ CD4+ (top panels ) and CD8+ (bottom panels), at baseline (T0), after the first dose (T1), 14 days after the second dose (T2), and 6 months after initial vaccination (T3). Time points were compared by non parametric repeated measures Friedman test; lines represent median with interquartile range. *p < 0.05; **p < 0.01; *** p < 0.001; ****p < 0.0001; no symbol, not significant. B) Fraction of AIM+ or total CD4+ (top panels) and CD8+ (bottom panels) that belong to the indicated subsets at each time point.

Thus, although the absolute number of antigen-specific T cells does not significantly increase following the second dose, the composition of this subset differs significantly between priming and recall responses, and indicates the establishment of an efficient population of S-specific cells comprising a significant fraction of non-terminally differentiated and potentially long-lived memory cells.

### Effector features of Spike-Specific CD4+ and CD8+ cells

We then explored by unbiased analysis the changes in time in the expression of immunologically relevant markers occurring in antigen-specific T cells. We used FlowSOM clustering to identify 4 CD4+ T cell clusters segregated by the expression pattern of CD137, CD39, ICOS, PD-1, HLA-DR, CD25, CXCR5, CD95, CCR7, CD45RA, CD38, and CD127 and superimposed those clusters in a UMAP plot generated by embedding the same set of markers (Fig. 5A). Since Ki67, a marker of cell proliferation, was never expressed on AIM+ cells (not shown), we excluded it from analysis. AIM+ CD4+ T cells distribute differently in the UMAP plot at the three different time points (Fig. 5B), highlighting an overall phenotypic shift. Among these cell clusters, cluster 3 and 4 result unchanged in frequency, while clusters 1 and 2 show significant changes over time (Fig. 5C). Cluster 2, which increases in frequency at each time point, is characterized by high expression of activation markers (CD25, CD38, CD39, HLA-DR, CD137) and displays features typical of T follicular helper (Tfh) cells (i.e. expression of ICOS-L, PD-1, and CXCR5) while cluster 1 is composed by cells which show a non-activated profile, and decreases with time (Fig. 5A and C). This data indicates that the antigen-specific CD4+ T cell phenotype is shaped and “perfectioned” by the second encounter with the antigen, and the acquired phenotype persists in the peripheral blood for at least 6 months.

**Fig.5:**
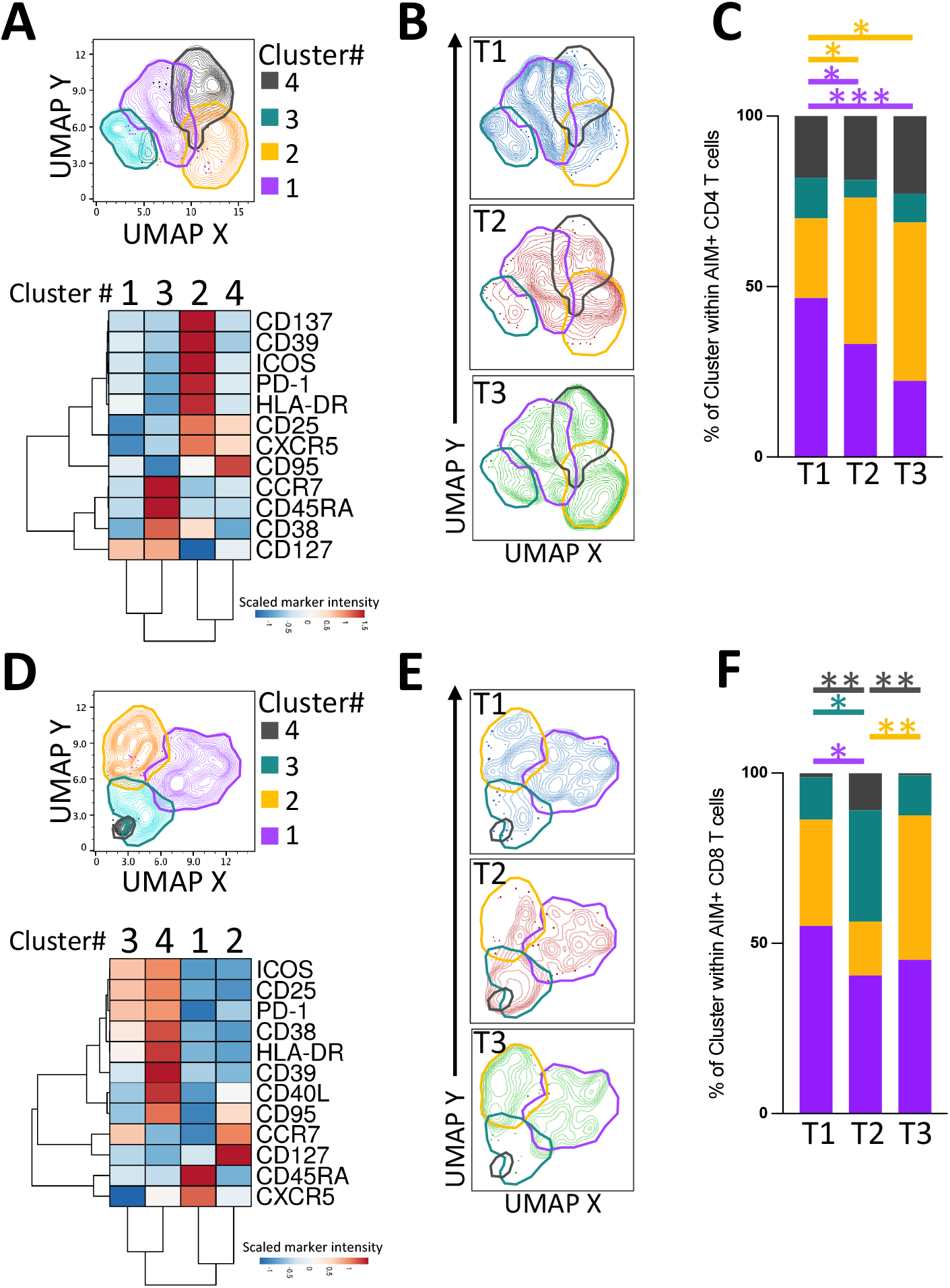
Phenotype shifts in AIM+ T cells over time. A) and D), top panels. UMAP embedding and FlowSOM clustering based on the expression of the indicated markers on AIM+ CD4 (A) and CD8 T cells (D). The cell clusters identified by FlowSOM are superimposed on the UMAP plots and manually contoured to highlight cluster boundaries. A and D) Bottom panels. Heatmaps with two-way hyerachical clustering of the scaled and centred median fluorescence intensity (MFI) values for the indicated markers expressed by the AIM+ CD4+ (A) and CD8+ (D) T cell clusters. B and E) UMAP plots with AIM+ CD4+ (B) and CD8+ (E) T cells selected from each time point. C and F) Bar plots showing relative frequency of cell clusters at indicated time points among AIM+ CD4+ (C) and CD8+ (F) T cells. Statistical significance was inferred by ANOVA (non parametric Friedman test) followed by a post hoc two-stage linear step-up procedure of Benjamini, Krieger and Yekutieli. *p < 0.05; **p < 0.01; *** p < 0.001.

Data analysis through manual gating and measurement of single marker expression on the antigen-specific T cells, revealed the emergence and persistence in time, albeit at lower levels after 6 months, of CD4+ subsets displaying high levels of PD-1, ICOS, and CXCR5 (Fig. S4), denoting their ability to interact with B cells in lymphoid follicles. At the 6-month time point expression of activation markers such as CD25, CD39, CD38, and HLA-DR is greatly reduced, in line with what expected after the physiological contraction of the immune response.

Also S-specific CD8+ lymphocytes are remodeled by vaccine doses, as shown by the distinct distribution with time along the UMAP axes (Fig. 5E). Two clusters, comprising highly (Cluster 4) and slightly less (Cluster 3) activated AIM+ CD8+ T cells, show transient expansion after the booster dose (Fig. 5 D and F). This is in contrast to Cluster 1 (Naïve-like cells) which remains stable after a first decrease in frequency at T2, and to Cluster 2 (CCR7+ CD127+ CD45RA- central memory CD8+ T cells) which is not significantly reduced at T2 but increases in size at 6-months. Manual analysis of AIM+ CD8+ T cells confirmed the transient increase at T2 of CD38+, HLA-DR+ and CD25+ cell frequencies, whereas CD39+ and PD-1+ cells steadily decrease or increase, respectively, at T2 and T3 compared to T1 (Fig. S4).

Thus, antigen-specific T cells acquire phenotypic features of activation and functional capability early after the booster, and most of these attributes are less evident and partially replaced, in the long run, by characteristics distinctive of more quiescent memory cells.

### Generation of T_SCM_ cells

Among the desirable outcomes of vaccination lies the generation of a pool of stem memory cells, which can rapidly and efficiently differentiate in an army of highly effective and polyfunctional lymphocytes in case of re-encounter with their nominal antigen *(31)*. Stemness includes long-term persistence, a key aspect in this age of pandemics and uncertainties on the durability of the novel vaccines. Thus, we searched for these cells within the antigen-responsive CD4+ and CD8+ subsets.

After the first dose, CD4+AIM+CD45RA+CD27+CCR7^high^CD95+ cells, representing CD4+T_SCM_ (T Stem cell memory) cells, were detectable in 88% of individuals (Fig. S5); 2 weeks after the second dose, this fraction was still 88%; after 6 months, 91% of individuals showed these cells (Fig.6A). The number of circulating CD4+ T_SCM_ rose from a median of 0,01 cells/ml at baseline to 14,7 cells/ml on the day of the boost, and further to 15,28 cell/ml two weeks after the second dose; after 6 months these cells were slightly increased in number (median 23,9 cells/ml). CD8+ T_SCM_ followed similar kinetics, with 94%, 87%, and 96% of individuals showing these cells (0.05, 6, 8, and 8 cells/ml at baseline, T1, T2, and T3 respectively). To investigate the possible impact of T_SCM_ on future immune responses, we applied a machine learning approach to test if the number of T_SCM_ induced by vaccination significantly predicts immunological outcomes at distant time points. We find that indeed the number of T_SCM_ induced after priming is a significant predictor of the number of both CD4+ and CD8+ activated T cells at the latest time point (Fig. 6B).

**Fig.6:**
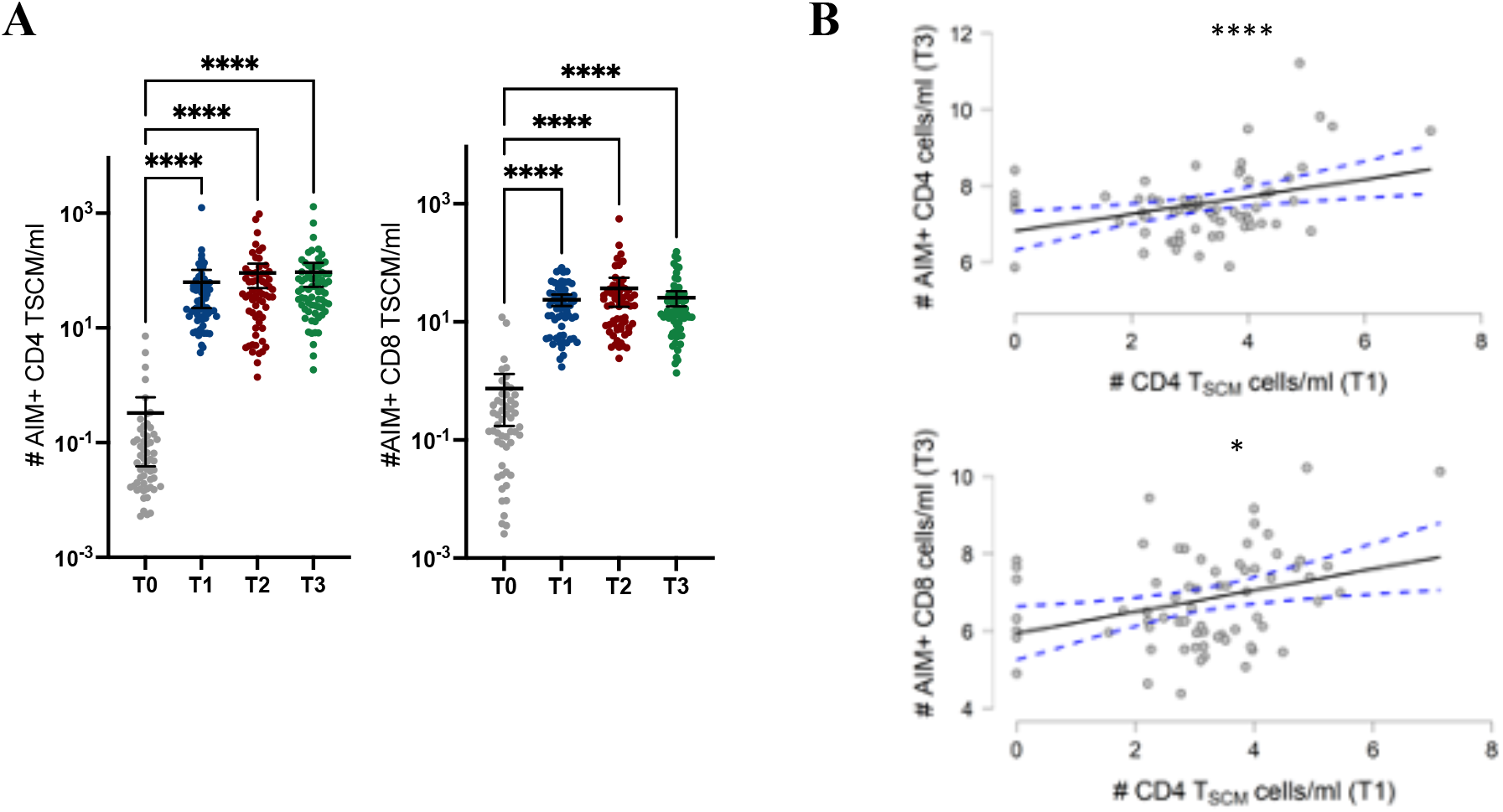
T_SCM_ cells are induced after priming and remain in the periphery for at least 6 months. A) Absolute cell counts of CCR7+CD45RA+CD27+CD95+ AIM+ CD4 (left) and CD8 (right) cells at the different time points. Timepoints were compared by non parametric Kruskall-Wallis test; lines represent median with interquartile range. ****p < 0.0001; no symbol, not significant. B) The relevance of T1 CD4 T_SCM_ cells/ml in predicting T3 AIM+ CD4 (top) and CD8 (bottom) cells/ml was tested with two General Linear Models with stepwise selection aiming to optimize Akaike Information Criterion (AIC) (Bozdogan, 1987). Model’s R2 were 0.21 and 0.16 for AIM+ CD4 and CD8 cells respectively. The number of CD4 T_SCM_ in T1 cells/ml was deemed as significant in predicting both CD4+ and CD8+ AIM+ cell numbers in T3 (****p < 0.0001; *p < 0.05). Lines represent linear regression.

These findings show that vaccination with BNT162b2 induces the emergence of a population of cells with features of longevity, which remain numerically stable in the peripheral blood for at least 6 months, and which predict the persistence of T cell responses.

## DISCUSSION

Mass vaccinations are underway to control the continued dissemination of SARS-CoV-2. Specific viral control is achieved through the action of effector cells of the adaptive immune system: the antibody-producing progeny of B cells, and the functionally diverse population of T cells derived from a small pool of naïve progenitors which expand and differentiate to achieve the ability to provide B cell help, to directly kill virally infected cells, and to sustain the immune response through the production of cytokines, all while maintaining a source of non-terminally differentiated but antigen-experienced cells which can rapidly expand upon antigen re-encounter. In all of this, T cells are indispensable: an optimal antibody response is the consequence of a competent underlying T cell response, and T cell responses alone can successfully clear infection with SARS-CoV-2, as shown in COVID-19 patients lacking B cells *(10, 12)*, and in seronegative COVID-19-recovered individuals *(32–34)*. Here, we investigate the immune response occurring after vaccination with BNT162b2 to understand how the vaccine animates T cells specific for the S-protein, in order to predict whether this response will be durable and if it is similar in individuals of both sexes and at different ages. The timing of our glimpses into the immune system’s antigen-specific dynamics was such that we could observe the emergence and the evolution of the effector functions of vaccine-induced T cells, and then record the physiological contraction of the immune response and study the features of the surviving antigen-specific cells. The use of freshly obtained blood cells permitted the detection of fragile markers, avoided the bias introduced by freezing/thawing procedures, and provided the possibility to precisely calculate absolute cell counts, a measure routinely used to guide clinical decisions in infectious diseases, such as HIV infection *(35)*.

The simultaneous measurement of serum levels of neutralizing antibodies provided insights on the effective induction of the humoral arm of the adaptive response. NAbs were induced by vaccination in all individuals in our cohort. Although titers did decline with time, they were maintained at high levels for the entire period of our observation (6 months), in agreement with previous studies *(27, 36)*. Importantly, the efficacy of the vaccine in inducing antibodies was equivalent in both sexes, and correlated inversely with age, as expected.

We find that most individuals harbour Spike-specific T cells already at baseline, likely due to the presence of a pool of naïve progenitors and of memory clones which are cross-reactive with other coronaviruses *(3, 15–17, 37, 38)*. These cells are highly responsive to antigen encounter, and their SI increases with time. CD8+ cells show a less vigorous response compared to the CD4+ subset possibly due to the sub-optimal stimulation of CD8+ cells by 15-mers, as those used in our assays *(39)*.

High antibody titers induced by influenza vaccination have been shown to correlate positively with the frequency of T cells expressing follicular helper molecules, including CXCR5 and ICOS *(40, 41)*. Relevant to infection with SARS-CoV-2, an interesting study has shown the absence of germinal centers and a block in the differentiation of Tfh cells in post-mortem tissues from COVID-19 patients *(42)*, with loss of both transitional and follicular B cells in severe disease. Here, we find a consistent population of T cells equipped with the molecules needed for interaction with B cells, which survives the contraction of the immune response and is clearly detectable 6 months after vaccination albeit with a measurable (but not statistically significant) decrease in magnitude. The number of antigen-specific cells expressing CXCR5, crucial for positioning T cells in the germinal center within lymph nodes, was increased 5-fold at the latest time point compared to the initial measurement on the emerging population of antigen-specific cells, three weeks after the first dose. Similarly, T cells expressing ICOS and PD-1 composed a significant fraction of the antigen-specific subset, and were numerically maintained for at least 6 months. These findings show that vaccination with BNT162b2 appropriately induces the differentiation of the T cell subset specialized in providing B cell help and thus in sustaining the generation of high-affinity antibodies in germinal centers, and that these cells persist in the periphery for at least 6 months. Importantly, AIM+ T cell numbers correlate with serum antibody levels only in the primary response, measured 3 weeks after the first dose, while subsequent measurements indicated that T cell responses and antibody levels follow different kinetics: thus, the sole measurement of nAbs does not inform on the whole immunological ensemble induced by vaccination, although high antibody levels seem to correlate with protection from infection both in animal models and in humans *(43–46)*.

Moreover, CD4+ T cells have also other abilities, such as those related to cytokine production and to direct cytotoxic function. In agreement with previous results, we find that antigen-specific CD4+ T cells induced by vaccination with BNT162b2 produce cytokines typical of the Th1 profile (IFNγ, TNFα), and minimal levels of IL-4 and IL-17. In infants the presence in the peripheral blood of ≥100 influenza-specific IFNγ-producing cells/ml confers protection against clinical influenza *(47)*, and a correlation between both magnitude and polyfunctionality of T cell responses and resistance to infection has been described in other vaccine settings *(48–54)*. In this study, we show that 6 months after vaccination with BNT162b2 over 300 CD4+ and CD8+ Spike-specific IFNγ-producing T cells/ml are clearly detected in the periphery, with diverse functionalities. CD4+ cells acquire both helper and cytotoxic functions which are particularly evident at the peak of the antigen-specific response, 14 days after the boost. At this time point we detect the highest frequencies of cells with a polyfunctional phenotype, in both CD4+ and CD8+ S-specific T cells. Interestingly, the CD4+ subset is more spared from the physiological contraction of the immune response, 6 months after receiving the first dose of BNT162b2, and it shows polyfunctional features of both cytotoxicity and ability to provide B-cell help. At the same time point, the antigen-specific CD8+ T cell subset seems to drop at lower levels, although these cells are still clearly detectable and are significantly expanded compared to the baseline. This data is consistent with studies which have investigated cellular immune responses in individuals recovering from COVID-19 *(16, 20, 55–58)*, and with other studies on mRNA vaccination *(59, 60)*, and suggest that vaccine-induced T cells have the prerequisites to confer protection from subsequent infections. Moreover, the persistence of a numerically consistent pool of antigen-specific T cell, which we find to be still increased 6-fold from baseline after 6 months, may be the source for of effector cells which can promptly expand in case of antigen re-encounter *(61)*.

The finding that CD4+ and CD8+ T_SCM_ are present throughout the 6-month period of this study suggests that immunity offered by vaccination should be long-lived, since it induces a reservoir of cells with multipotent capacity which likely are the very cells that provide long-lasting protection, and whose generation is a target of vaccination *(62–64)*. Importantly, the number of T_SCM_ was stable during the period of our study, and crucially those induced after priming are highly significant predictors of future T cell responses. This knowledge may inform current vaccination strategies and decisions on third dose vaccine administration, which may be spared for the fragile or immunologically impaired and re-directed to the unvaccinated, globally.

This study has the inevitable limits of human studies, and was performed on circulating lymphocytes, which may be different from those who reside at mucosal barriers and which confer immediate protection against infection. Also, we did not investigate Spike-specific B lymphocytes, and dosage of antibody titers was the only read-out of successful induction of humoral immunity. Similarly, the innate arm of the immune response, which has intriguingly been shown to be activated by vaccination*(65)*, was not included in this study; both of these elements are currently under active investigation the world over. Further time points will also be necessary to measure effective durability of anti-SARS-CoV-2 immune responses. Moreover, although donors were equally distributed for age and sex, our sample was limited in size.

This notwithstanding, some considerations may be made. On the whole, the results of this study can be visualized as the dynamic and integrated emergence of a theoretically effective Spike-specific adaptive immune response, characterized by a T cell response which precedes in time the development of high levels of anti-Spike neutralizing antibodies (Fig.7). T cells induced after the first encounter with the Spike protein are mostly effector cells, which secrete intermediate levels of cytokines, express the highest levels of Granzyme B, have not acquired polyfunctionality, and a fraction is also competent for the interaction with B cells; notably, also cells with features of stemness and longevity appear. After the boost, the peak of the response shows a fully activated, cytotoxically empowered, multifunctional, and B cell “friendly” T cell population, and this also corresponds to the highest levels of neutralizing antibodies in the serum. What remains after 6 months is a population of T cells with features of polyfunctionality and ability to provide B cell help; at this time point, Spike-specific T cells which have survived the immunological contraction are highly specific and prone to give rise to effective and rapid antiviral responses, both by sustaining the production of neutralizing antibodies and by exerting direct cytotoxicity towards already infected cells. Concomitantly, the pool of T_SCM_ is stably maintained. In our cohort we did not find differences between males and females in any of the investigated immunological measurements.

**Fig.7:**
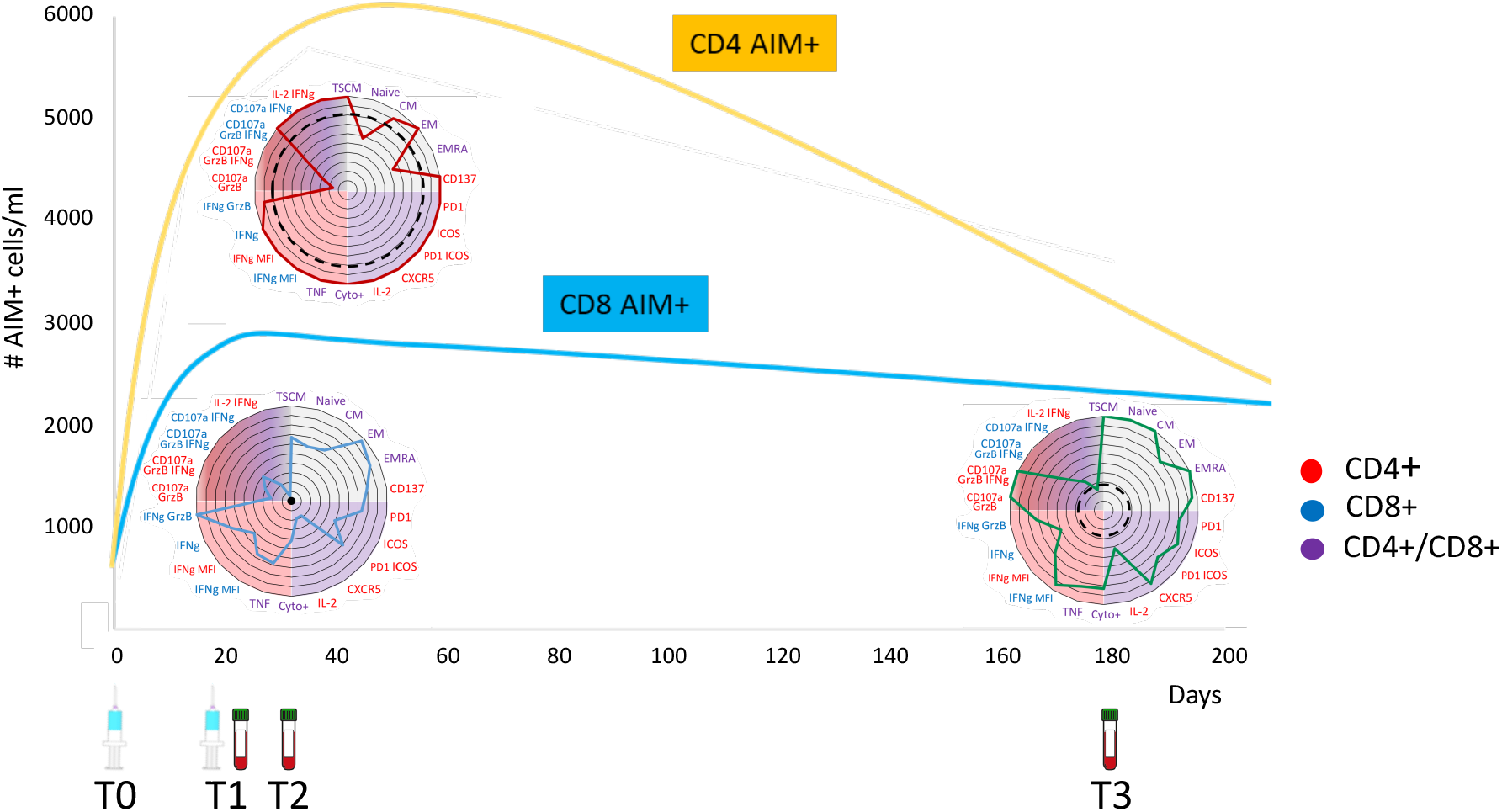
T cell responses to vaccination with BNTb16b2. T cell marker measurements from both CD4 and CD8 AIM+ cells are normalized across the 3 time points, and the relative positivity for each marker is displayed on the radar plots. Measurements from CD4+ T cells are shown in red, those from CD8+ cells are blue, and cumulative measurements from both subsets are purple. The resulting plots illustrate the main features of T cells responding to a pool of peptides derived from the Spike protein of SARS-CoV-2. Dashed circles indicate neutralizing anti-Spike antibody levels. Histograms represent CD4 (yellow) and CD8 (blue) cell counts along the timeline. Siringes indicate the time point of vaccine administration, and tubes correspond to the day of blood sampling. T_SCM_: T memory stem cells; CM: central memory; EM: effector memory; EMRA: CD45RA+ effector memory; Cyto+: aggregation of absolute numbers of CD4 and CD8 cell producing at least one cytokine among IFNγ, IL2, and TNF-α IFNγ MFI: mean fluorescence intensity of the IFNγ signal in IFNγ+ CD8 (blue) or CD4 (red) cells.

This model provides the immunological basis of the current clinical data showing the effectiveness of the vaccine in preventing severe disease, of the low frequency of breakthrough infections, and of the reduced time-window of contagiousness of re-infected individuals, all signs of effective immunity. Previous literature on T cell responses following vaccination shows similar immunological patterns in previously validated and approved anti-viral vaccines with outcomes of pluri-decennial immunity. This also suggests that a third booster dose may not be needed in most individuals, since over 90% of vaccinees develop T cells with features of longevity and show high numbers of circulating and appropriately modeled antigen-specific memory cells, ready to effectively and rapidly engage in a possible re-encounter with the virus. Immunologically fragile individuals, however, or those who are unavoidably exposed to high titers of virus may be better protected by a third dose of vaccination which has been shown to increase antibody levels, thus providing an immediate albeit temporary shield against viral entry in the host cell. In an equitable world, and based on the current data on vaccine effectiveness in preventing severe disease, after having secured the fragile from infection the absolute precedence should be given to unvaccinated individuals globally, in the unified endeavor to curb viral circulation and to prevent disease.

## MATERIALS AND METHODS

### Study Design

This work started to prospectively define cellular immune responses against the SARS-CoV-2 Spike-protein induced by mRNA vaccine administration. To this aim, we enrolled 71 individuals from the sanitary personnel and from scientists operating at the Santa Lucia Foundation which were scheduled for vaccination with Pfizer-BioNTech BNT162b2 between 12 January and 2 February 2021 (Table S1). All donors signed informed consent forms approved by the Ethical Committee of the Santa Lucia Foundation. Venous blood was obtained immediately before the first dose (T0), 21 days thereafter, at the time of boosting (T1), and two weeks after the second dose (T2), and 6 months after the first dose (T3). To avoid gender and age biases, an equal number of female and male volunteers, as well as an equal number of subjects 22-45 and 45-66 years old, was included. Not all subjects were analyzed at each time point due to precautionary quarantine of some individuals during the study. All data is presented in table 3.

### Evaluation of anti SARS-CoV-2 Antibodies

The measurement of anti SARS-CoV-2 neutralizing Abs was performed by electrochemiluminescence sandwich immunoassay (ECLIA) through Roche Elecsys Anti-SARS-CoV-2 S (Roche diagnostics, Switzerland). The neutralizing Ab were measured on a Cobas 601 modular analyzer (Roche diagnostics, Switzerland), using a cut-off of 0.8 U/ml to determine Ab levels. In particular, Elecsys Anti-SARS-CoV-2 S U/mL measurements are equivalent to WHO International Standard Binding Arbitrary Units per mL (BAU/mL), according to which higher values than 0.8 BAU/mL are considered positive.

### AIM assay

In vitro stimulation of freshly obtained PBMC was performed as described *(66)*. In brief, 200 ml of cell suspensions (10 x 10^6^ cells/ml) were seeded in U-bottom 96-well plates at a density of (0,2 ml/well) and stimulated for 18 hours with or without PepTivator SARS-CoV-2 protein S, S1 and S+ peptide pools (1 mg/ml each, Miltenyi Biotec). Purified αCD40 (0,5 μg/μl, Miltenyi Biotech) was added at culture start to enhance endpoint CD40L staining by inhibiting its recycling. Supernatants and cells from these culture wells were collected for cytokine measurement and flow cytometry, respectively.

### Intracellular Cytokine Staining

PBMC were incubated with the addition of BV421-conjugated antiCD107a (1/200 dilution, BD Biosciences), Monensin and Brefeldin A (5μM and 10 μg/ml, respectively, both from Sigma-Aldrich) after the first hour of incubation to allow accumulation of cytokines into the cytoplasm and avoid endo-lysosome acidification which can quench fluorescence from the αCD107a reinternalized during degranulation. At the end of the culture, cells were harvested and directly stained for flow cytometry.

### Flow cytometry staining and acquisition

Post-culture cells were pelleted in V-bottom 96-well plates and resuspended in 30 μl of antibodies at pre-optimized concentrations and diluted in Brilliant Stain Buffer (BD Biosciences), then incubated for 15’ at RT. After a washing step, cell pellets were fixed either in FoxP3 fixation/permeabilization Buffer (ThermoFisher) for 20’ at RT (AIM Assay) or in Formaldehyde 4% in PBS (ICS Assay). ICS was performed by incubating cells in 30 μl of antibodies against cytokines and Granzyme B diluted in a Saponin (Sigma-Aldrich) 0.5% W/V solution. The complete list of antibodies used for surface staining and ICS is shown in Table S2. Samples were acquired on a fully equipped Cytoflex LX within 6 hours from staining and fixation. Quality Control beads (Beckman Coulter) were used daily to check and standardize instrument performances. Data was analyzed with FlowJo v. 10.7

### Cytokine measurements

IL4, IL17, TNFα and IFNγ were measured in frozen culture supernatants from the AIM assays described above by a bead-based sandwich ELISA (MACSPlex Cytokine Kit, Miltenyi Biotech) according to manufacturer instructions.

### Statistical analysis

Statistical analysis was performed with GraphPad Prism 9. Details of the statistical tests applied for each experiment are illustrated in the respective figure legends.

## Supplementary Materials

Fig. S1 Identification of AIM+ and IFNγ+ T cell subpopulations by flow cytometry.

Fig. S2 Correlation of T cell responses with age and sex.

Fig. S3 Cytokine production by Spike-specific T cells.

Fig. S4 Phenotype of AIM+ CD4+ and CD8+ AIM+ T cells

Fig. S5 Vaccination with BNT162b2 induces T cells with features of T_SCM_.

Table S1 Donors

Table S2 Flow Cytometry Reagents

## Acknowledgments

We thank the volunteers for donating their blood and time, and the nurses for their assistance. We are grateful to Daniela F. Angelini for critically reading the manuscript.

## Funding

This work was partially supported by the Italian Ministry of Health to L.B. (COVID-2020-12371735).

## Author contributions

GB, LB, and AR conceptualized the study. AR, AS and CC performed vaccination supervision and donor enrollment. GG and MP set up and supervised all experimental procedures. GG, RP, SD, AV, MPirr, MS, FG performed cell stimulations, stainings, flow cytometry experiments, and cytokine measurements. MPB performed antibody dosage. GB prepared the dataset for analysis. GB and MP performed descriptive statistics. AT and CF did the statistical analysis. Funding acquisition: LB. Writing – original draft: GB. Writing – review & editing: GB, MP, LB, with input from all authors.

## Competing interests

The authors declare that they have no competing interests.

## Data and materials availability

All data is included in Table S3.

## Notes

### Competing Interest Statement

The authors have declared no competing interest.

