## Supplementary figures for "The BNT162b2 mRNA vaccine induces polyfunctional T cell responses with features of longevity"

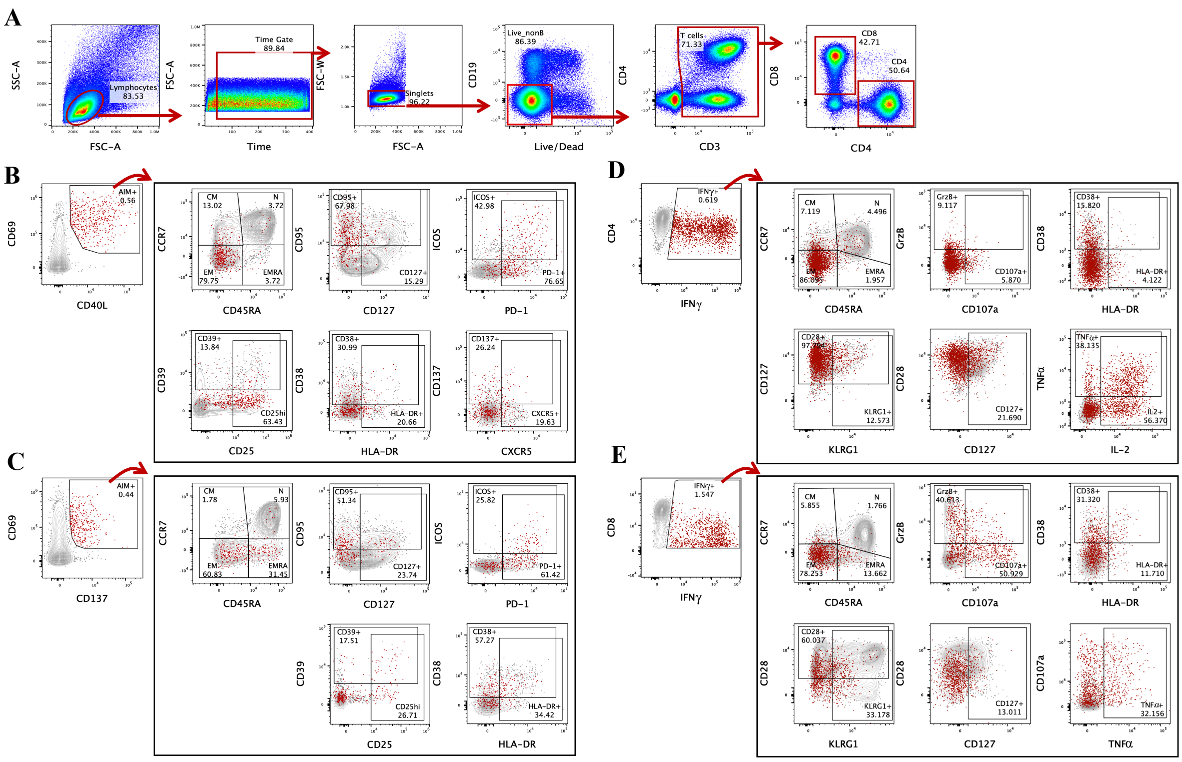

**Fig.S1 Identification of AIM+ and IFNγ+ T cell subpopulations** A) Gating of viable single T cells. A common gating is applied, which sequentially selects for lymphocytes by scatter analysis followed by a Time Gate used to exclude fluidics drifts during sample acquisition, for live non-B cells by excluding CD19+ and Live/Dead+ events and for CD3+ CD4+/CD8+ cells by conventional bivariate plotting. B-E) Gating of AIM+ (B and C) and IFNγ+ (D and E) CD4 (B and D) and CD8 (C and E) T cells. Red dots represent AIM+ or IFNγ+ cells overlaid on total CD4+ or CD8+ cells (in gray), while numbers inside or adjacent to the gates indicate frequencies among parent populations.

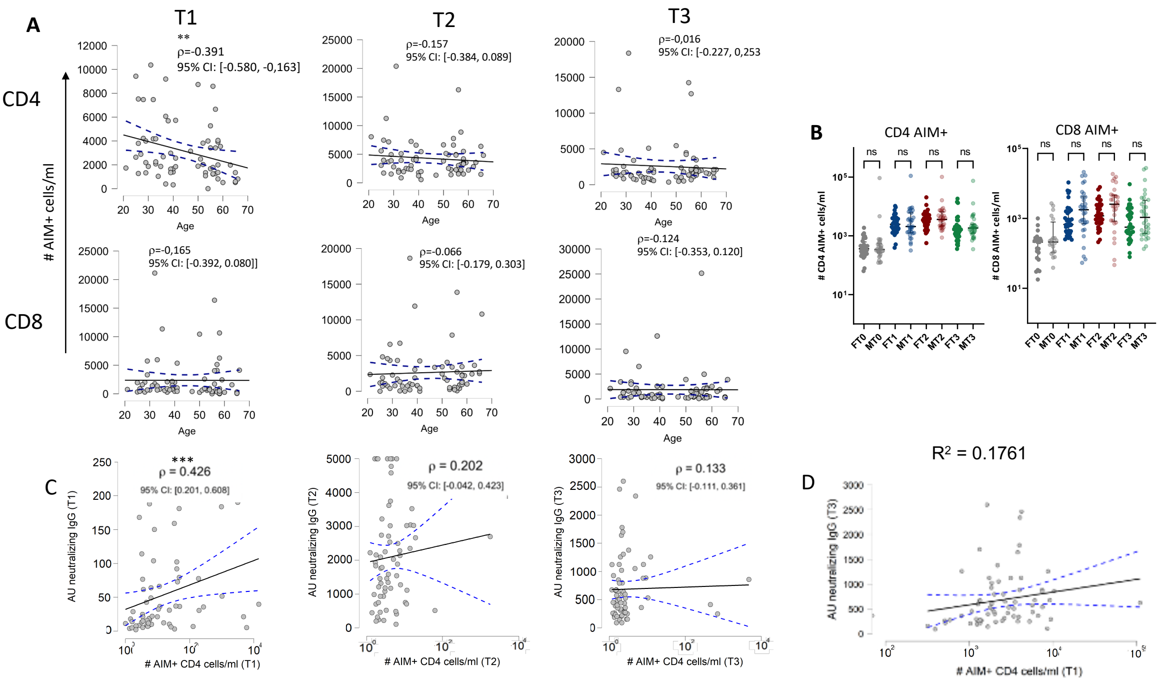

**Fig. S2 Correlation of T cell responses with age and sex.** A) Correlation plots of absolute CD4 (top) and CD8 (bottom) T cell counts and age at the different time points: 3 weeks after the first dose (T1) , 2 weeks after the second dose (T2), and 6 months after initial vaccination. Lines represent linear regression. Spearman's rank correlation test was used to measure significance **=p<0,01. B) Absolute AIM+ CD4 (top) and CD8 (bottom) T cell counts in females (F) and males (M) at each time point. Kruskall-Wallis test with Dunn correction for multiple comparisons was used to test significance; ns= not significant. C) Spearman's rank correlation test was applied to test for correlation between AU neutralizing IgG and the number of AIM+ CD4 cells/ml at each time point (T). Significance levels, Spearman's rank correlation coefficient (ρ) and Confidence Intervals for each plot are reported in figure (* = p < 0.05; ** = p < 0.01; *** = p < 0.001; **** = p < 0.0001). D) A linear regression model was fitted to test for T1 significant predictors of T3 AU neutralizing IgG. Variables were log10-scaled when their distribution wasn't normal. Model's $R^2$ was 0.18. The number of AIM+ CD4 cells/ml was deemed as significant **p < 0.01. Lines represent linear regression.

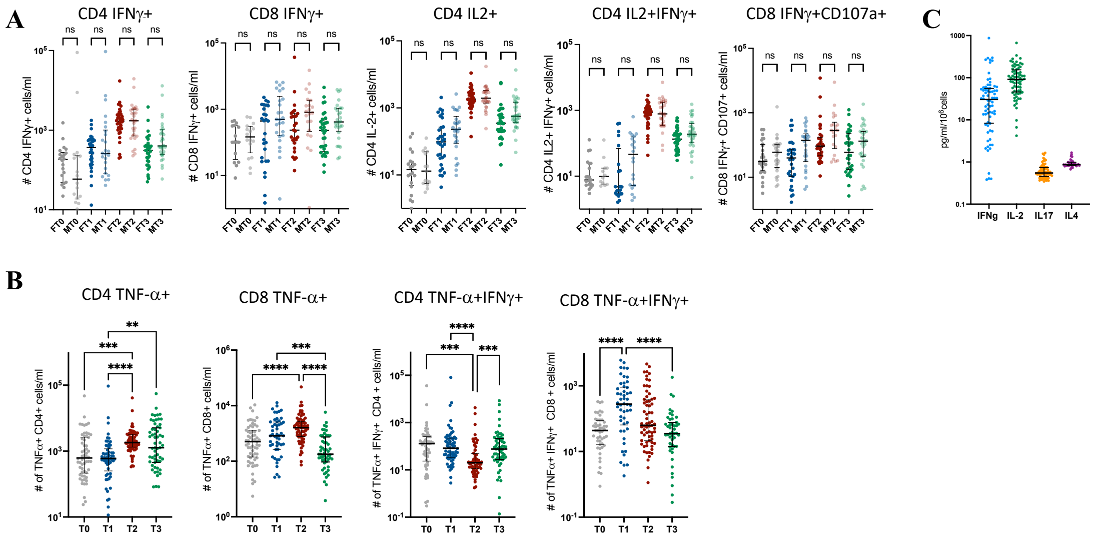

**Fig. S3 Cytokine production by Spike-specific T cells.** A) Absolute cell counts of CD4 and CD8 cells producing cytokines in females and males following o.n. stimulation with a pool of overlapping peptides covering the wt Spike protein at baseline (T0), 21 days after the first dose (T1), 14 days after the second dose (T2), and 6 months after initial vaccination (T3), as determined by intracellular staining and flow cytometric analysis. Time points were compared by non parametric Kruskall-Wallis test; ns=not significant. B) Absolute cell counts of CD4 and CD8 cells producing TNFα or both TNFα and IFNγ. Time points were compared by non parametric repeated measures Friedman test; lines represent median with interquartile range. *p < 0.05; **p < 0.01; *** p < 0.001; ****p < 0,0001; no symbol, not significant. C) Cytokine levels in supernatants from o.n. cultures of PBMCs stimulated with Spike-protein peptides. Cells were obtained 2 weeks after the second dose (T2), and cytokines were measured using the MACSPlex assay.

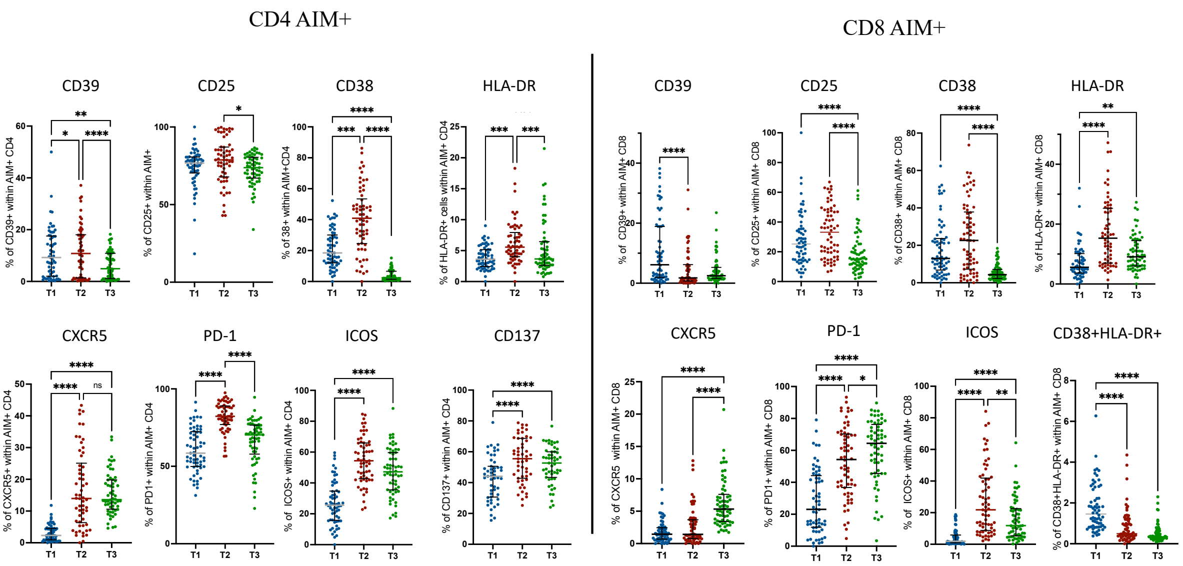

**Fig. S4 Phenotype of AIM+ CD4 and CD8 AIM+ T cells, at the different timepoints.** Timepoints were compared by non parametric repeated measures Friedman test; lines represent median with interquartile range. *p < 0.05; **p < 0.01; *** p < 0.001;  ****p < 0.0001; no symbol, not significant.

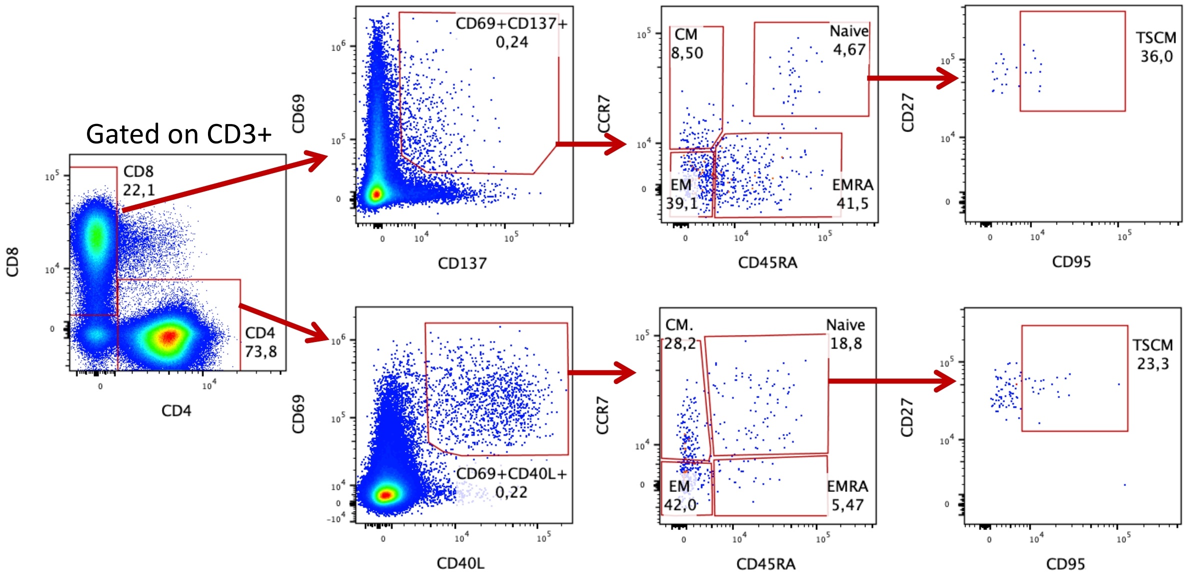

**Fig.S5: Vaccination with BNTb162b2 induces T cells with features of $T_{SCM}$** Flow cytometry plots from one representative female donor.

| **Table S1** | **Female** | **Male** | **All** |
|---|---|---|---|
| Number | 37 | 34 | 71 |
| Age (mean/range) | 44.2/25-65 | 45.2/21-66 | 47.7/21-66 |
| Race/Ethnicity | All caucasian | All caucasian | All caucasian |

| **Table S2** | | | | |
|---|---|---|---|---|
| | | **MIX AIM** | | |
| **Antibody** | **Fluorochrome** | **Clone** | **Company** | **Dilution** |
| CD3 | BUV496 | UCHT1 | Bect.Dick. | 1:100 |
| CD4 | iFluor810 | RPA-T4 | AAT Bioquest | 1:200 |
| CD8 | iFluor594 | SK1 | AAT Bioquest | 1:150 |
| CD19 | BV650 | SJ25C1 | Bect. Dick. | 1:40 |
| CD25 | BUV661 | 2A3 | Bect. Dick. | 1:60 |
| CD27 | PE-CF594 | M-T271 | Bect.Dick. | 1:300 |
| CD38 | PE-Cy5.5 | LS198-4-3 | Coulter | 1:100 |
| CD39 | BV605 | TU66 | Bect.Dick. | 1:30 |
| CD45RA | BV480 | HI100 | Bect. Dick. | 1:200 |
| CD69 | BB700 | FN50 | Bect. Dick. | 1:100 |
| CD95 | BB515 | DX2 | Bect. Dick. | 1:60 |
| CD127 | PE-Cy5 | R34.34 | Coulter | 1:150 |
| CD137 | BUV395 | 4B4-1 | Bect. Dick. | 1:30 |
| CD154(CD40L) | PE | 24-31 | eBioscience | 1:30 |
| CD185 (CXCR5) | APC-R700 | RF8B2 | Bect. Dick. | 1:60 |
| CD278 (ICOS) | APC | ISA-3 | Coulter | 1:100 |
| CD279 (PD-1) | BV421 | EH12.1 | Bect. Dick. | 1:50 |
| CCR7 | PE-Cy7 | G043H7 | SONY | 1:30 |
| HLA-DR | APC-Vio770 | REA805 | Miltenyi | 1:120 |
| Ki67 | BV785 | B56 | Bect. Dick. | 1:30 |
| Live Dead | Promo Fluor 840 | | Promokine | 1:10.000 |
| | | **MIX ICS** | | |
| **Antibody** | **Fluorochrome** | **Clone** | **Company** | **Dilution** |
| CD3 | BUV496 | UCHT1 | Bect.Dick. | 1:100 |
| CD4 | iFluor810 | RPA-T4 | AAT Bioquest | 1:200 |
| CD8 | iFluor594 | SK1 | AAT Bioquest | 1:150 |
| CD25 | BUV661 | 2A3 | Bect. Dick. | 1:60 |
| CD28 | APC-Vio770 | REA612 | Miltenyi | 1:120 |
| CD38 | PE-Cy5.5 | LS198-4-3 | Coulter | 1:100 |
| CD45RA | BV480 | HI100 | Bect. Dick. | 1:200 |
| CD107a | BV421 | H4A3 | Bect. Dick. | 1ul/well 10^5cells |
| CD127 | PE-Cy5 | R34.34 | Coulter | 1:150 |
| HLA-DR | BV785 | G46-6 | Bect. Dick. | 1:100 |
| TNF$\alpha$ | BV650 | MAb11 | Bect. Dick. | 1:30 |
| IL-2 | PE-CF594 | 5344.111 | Bect. Dick. | 1:50 |
| IFN$\gamma$ | APC | B27 | Bect. Dick. | 1:100 |
| GRZB | Alexa 700 | GB11 | Bect. Dick. | 1:100 |
| CCR7 | PE-Cy7 | G043H7 | SONY | 1:30 |
| KLRG1 | FITC | REA261 | Miltenyi | 1:100 |
| Live Dead | Promo Fluor 840 | | Promokine | 1:10.000 |
